# High-throughput and high-efficiency sample preparation for single-cell proteomics using a nested nanowell chip

**DOI:** 10.1101/2021.02.17.431689

**Authors:** Jongmin Woo, Sarah M. Williams, Victor Aguilera-Vazquez, Ryan L. Sontag, Ronald J. Moore, Lye Meng Markillie, Hardeep S. Mehta, Joshua Cantlon, Joshua N. Adkins, Richard D. Smith, Geremy C. Clair, Ljiljana Pasa-Tolic, Ying Zhu

## Abstract

Global quantification of protein abundances in single cells would provide more direct information on cellular function phenotypes and complement transcriptomics measurements. However, single-cell proteomics (scProteomics) is still immature and confronts technical challenges, including limited proteome coverage, poor reproducibility, as well as low throughput. Here we describe a nested nanoPOTS (N2) chip to dramatically improve protein recovery, operation robustness, and processing throughput for isobaric-labeling-based scProteomics workflow. The N2 chip allows reducing cell digestion volume to <30 nL and increasing processing capacity to > 240 single cells in one microchip. In the analysis of ∼100 individual cells from three different cell lines, we demonstrate the N2 chip-based scProteomics platform can robustly quantify ∼1500 proteins and reveal functional differences. Our analysis also reveals low protein abundance variations (median CVs < 16.3%), highlighting the utility of such measurements, and also suggesting the single-cell proteome is highly stable for the cells cultured under identical conditions.

## Introduction

With the success of single-cell genomics and transcriptomics, there is a growing demand for high-throughput single-cell proteomics (scProteomics) technologies. Global profiling of protein expressions in individual cells can potentially reveal specific protein markers accounting for heterogeneous populations, provide more concrete evidence of cellular function phenotypes, and help to identify critical post-translational modifications that regulate protein activities ^1-3^. Despite this transformative potentials, scProteomics still lags behind single-cell transcriptomics in terms of coverage, measurement throughput, and quantitation accuracy.^4^

Most reported mass-spectrometry-based scProteomics technologies can be classified based upon whether they make use of isotopic labeling; i.e. they are either label free or use isobaric labeling. In the label-free methods ^5-10^, single cells are individually processed and analyzed using liquid chromatography-mass spectrometry (LC-MS) signal intensity measurements (i.e. MS1 ion currents) to quantify protein abundance. To improve proteome coverage, high-recovery sample preparation systems ^9-11^ and highly sensitive LC-MS systems ^7, 12, 13^ are usually employed. Although label-free approaches exhibit better quantification accuracy and higher dynamic range, their throughputs are limited, as each cell requires > 0.5 hour-long LC-MS analysis. In the isobaric labeling approaches (e. g., tandem mass tags or TMT) ^14-18^, single-cell digests are labeled with unique isobaric labels, that are then pooled together for a multiplex LC-MS analysis. Importantly, the peptides originating from different single cells appear as a single MS1 peak. As a consequence, the pooled ions contributing to a given precursor peak is higher than from individual cells and their fragmentations result in a richer MS2 spectrum for peptide identification. The released reporter ions infer protein abundance in different single cells. A “carrier” sample containing a larger amount of peptides than individual cells (e.g. ∼100×) is spiked into each isobaric labeling pool to maximize the peptide identification (SCoPE-MS) ^14, 15, 18^. Currently, the isobaric-labeling approaches have enabled to analyze ∼100 single cells per day. We anticipate the throughput will increase gradually with new releases of higher multiplex isobaric reagents, shorter LC gradients, and the inclusion of ion mobility in single-cell proteomics pipelines.

Analogous to single-cell transcriptomics, microfluidic technologies play increasing roles in sample preparation for scProteomics ^6, 9, 11^. By minimizing the sample processing volumes in nanowells or droplets, the non-specific-binding-related protein/peptide loss is reduced, resulting in improved sample recovery. More importantly, both protein and enzyme concentrations increase in nanoliter volumes, enhancing tryptic digestion efficiency. For example, our lab developed a nanoPOTS (nanodroplet processing in one-pot for trace samples) platform for significantly improving proteomics sensitivity by minimizing the reaction volume to < 200 nL ^11^. NanoPOTS allowed reliably identifying 600–1000 proteins with label-free approaches ^7, 12, 13^. When isobaric labeling approaches were used, ∼1500 proteins could be quantified across 152 single cells and at a throughput of 77 per day ^6, 15^. Despite this progress, challenges remain. In current microfluidic approaches, the sample processing volume is >10,000 larger than a single cell and gains would be expected from further miniaturizing the volumes, but it is presently constrained by liquid handling operations, including reagent dispensing, sample aspirating, transferring, and combination. Among these, the nanoliter-scale aspirating and transferring steps, which are commonly performed in isobaric-labeling workflows, are challenging, time-consuming, and prone to sample losses. Additionally, most reported microfluidic approaches employed home-built nanoliter liquid handling systems, which limits their broad dissemination.

Herein, we describe a nested nanoPOTS (N2) chip to improve isobaric-labeling-based scProteomics workflow. Compared with our previous nanoPOTS chip, ^6, 15^ where nanowells are sparsely distributed, we cluster arrays of nanowells in dense areas and use them for digesting and labeling single cells with single TMT sets. With the N2 chip, we eliminate the tedious and time-consuming TMT pooling steps. Instead, the single-cell samples in one TMT set are pooled by simply adding a microliter droplet on top of the nested nanowell area and retrieving it for LC-MS analysis. The N2 chip reduces the sample processing volumes by one order of magnitude and allows over 5× more numbers of nanowells in one microchip for high-throughput single-cell preparation. We demonstrate the N2 chip not only efficiently streamlines the scProteomics workflow, but also dramatically improves sensitivity and reproducibility.

## Methods

### Fabrication and assembly of the N2 chips

The chips were fabricated on glass slides using standard photolithography, wet etching, and silane treatment approach as described previously ^11, 19^. Briefly, as shown in Figure 1 and S1a, 27 (3 × 9) nanowell clusters with a distance of 4.5 mm between adjacent clusters are designed on single microscope slide (1 × 3 inch, Telic Company, Valencia, USA). In each cluster, 9 nanowells with 0.5-mm diameter and 0.75-mm well-to-well distance are nested together. To facilitate droplet combination and retrieval process, a micro-ring surrounds the nested nanowells. After photoresist exposure, development, and chromium etching, the glass slide was etched to a depth of ∼5 µm with buffered hydrofluoric acid ^20^. The freshly etched slide was dried by heating it at 120 °C for 2 h and then treated with oxygen plasma for 3 min (AP-300, Nordson March, Concord, USA). To selectively pattern the chip, 2% (v/v) heptadecafluoro 1,1,2,2-tetrahydrodecyl-dimethylchlorosilane (PFDS, Gelest, Germany) in 2,2,4-trimethylpentane was applied on the chip surface and incubate for 30 min. After removing the remaining chromium layer, all the chromium-covered regions are (nanowells and micro-rings) are hydrophilic and exposed areas are hydrophobic. Finally, a glass frame was attached to the nanowell chip with epoxy to create a headspace for reaction incubation.

**Figure 1.**
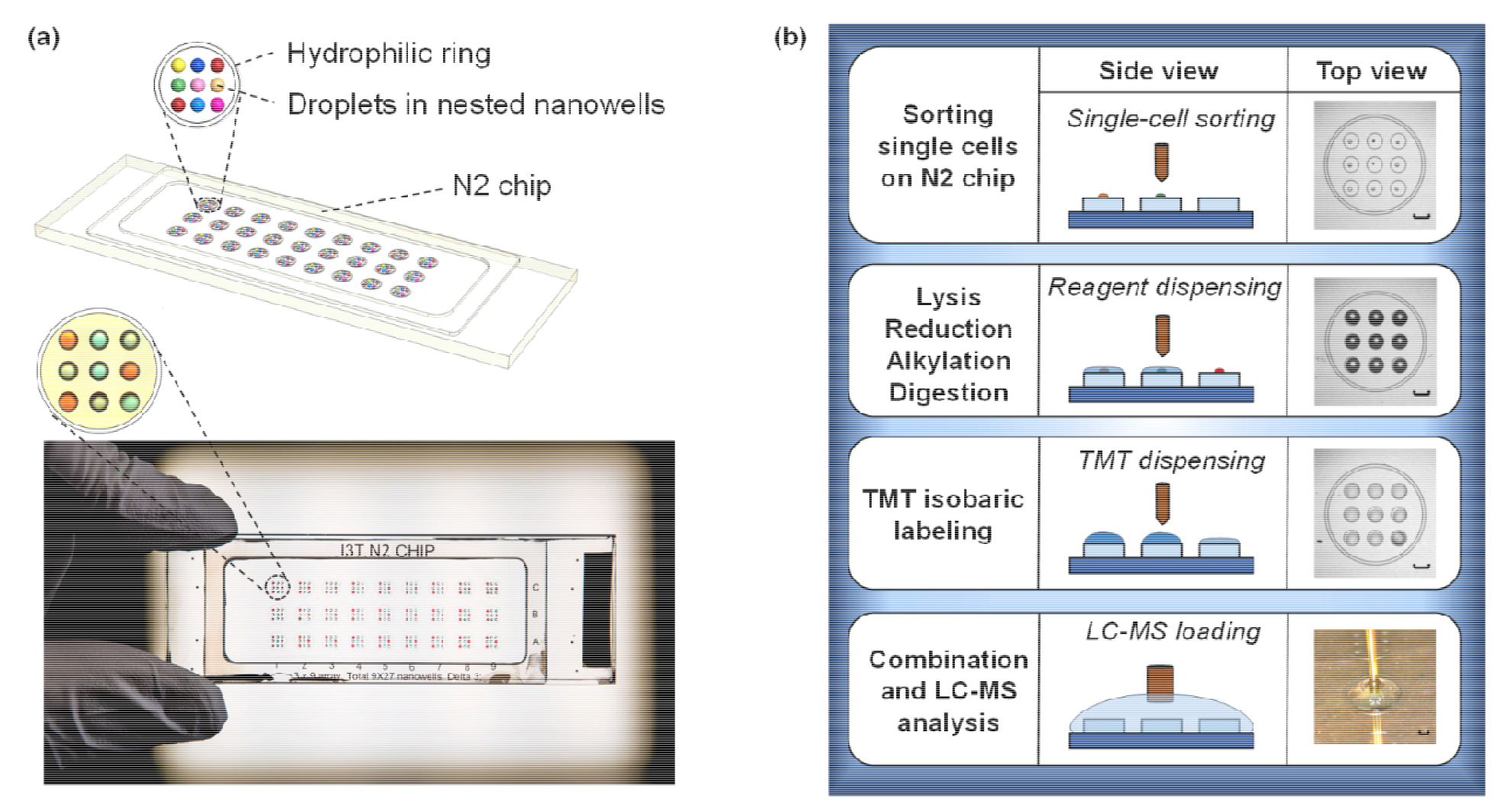
(a) A 3D illustration (top) and a photo (bottom) of the nested nanoPOTS (N2) chip. Nine nanowells are nested together and surrounded by a hydrophilic ring for an TMT set. (b) Single-cell proteomics workflow using the N2 chip. The scale bar is 0.5 mm.

### Reagents and chemicals

Urea, n-dodecyl-β-D-maltoside (DDM), Tris 2-carboxyethyl phosphine (TCEP), Iodoacetamide (IAA), Ammonium Bicarbonate (ABC), Triethylammonium bicarbonate (TEAB), Trifluoroacetic acid (TFA), Anhydrous acetonitrile (a-ACN), and Formic acid (FA) were obtained from Sigma (St. Louis, MO, USA). Trypsin (Promega, Madison, WI, USA) and Lys-C (Wako, Japan) were dissolved in 100 mM TEAB before usage. TMTpro 16plex, 50% hydroxylamine (HA), Calcein AM, Acetonitrile (ACN) with 0.1% of FA, and Water with 0.1% of FA (MS grade) were purchased from Thermo Fisher Scientific (Waltham, MA, USA).

### Cell culture

Three murine cell lines (RAW 264.7, a macrophage cell line; C10, a respiratory epithelial cell line; SVEC, an endothelial cell line) were obtained from ATCC and cultured at 37°C and 5% CO2 in Dulbecco’s Modified Eagle’s Medium supplemented with 10% fetal bovine serum and 1× penicillin-streptomycin (Sigma, St. Louis, MO, USA).

### Bulk-scale proteomic sample preparation and mimic single-cell experiments

The cultured cell lines were collected in a 15 ml tube and centrifuged at 1,000 × g for 3 min to remove the medium. Cell pellets were washed three times by 1× PBS buffer, then counted to obtain cell concentration. Ten million cells per cell population were lysed in lysis buffer containing 8M urea in 50 mM ABC in ice. Protein concentration was measured with BCA assay. After protein was reduced and alkylated by DTT and IAA, Lys-C (enzyme-to-protein ratio of 1:40) was added and incubated for 4 h at 37°C. Trypsin (enzyme-to-protein ratio of 1:20) was added and incubated overnight at 37°C. The digested tryptic peptides were acidified with 0.1 % TFA, desalted by C18 SPE column, and completely dried to remove the acidic buffer.

After measuring the peptide concentration with BCA assay, samples from three different cell types were mixed at 1:1:1 ratio and used for boost and reference samples. All peptide samples were dissolved with 50 mM HEPES (pH 8.5) followed by mixing with a TMT-16 reagent in 100% ACN. To maintain high labeling efficiency, a TMT-to-peptide ratio of 1:4 (w/w) was used. After 1-h incubation at room temperature, the labeling reaction was terminated by adding 5% HA and incubating for 15 min. The TMT-labeled peptides were then acidified with 0.1% FA and cleaned with C18 stage tips. Before use, different amounts of peptides (0.1 ng for mimic single cell, 0.5 ng for reference, 10 ng for boost) were diluted in 0.1% FA buffer containing 0.1% DDM (w/v) to prevent sample loss at low concentration conditions.

To mimic single-cell proteomics preparation in nanowell chips, 0.1-ng peptide samples in 200 nL buffer from the three cell lines were loaded into 1.2-mm nanowells using a nanoPOTS dispensing robot ^11^ and incubated for 2 h at room temperature. Next, samples from the same TMT set were collected and combined into a large-size microwell (2.2-mm diameter), which contained 10 ng and 0.5 ng TMT-labeled peptides for boost and reference samples, respectively.

To deposit these single-cell-level peptide samples to N2 chip, we employ a picoliter dispensing system (cellenONE F1.4, Cellenion, France) to dispense 0.1-ng peptide in 20 nL buffer in each nanowells (Figure S1b). After incubating the chip at room temperature for 2 h, mixed boost and reference samples (10 ng and 0.5 ng, respectively) were equally distributed into each nanowell.

Samples in both nanowell chip and N2 chip were completely dried out in a vacuum desiccator and stored in a -20°C freezer until analysis.

### scProteomics sample preparation using the N2 Chip

The cellenONE system was used for both single-cell sorting and sample preparation on N2 chip. Before cell sorting, all the cells were labeled with Calcein AM (Thermo Fisher) to gate out dead cells and cell debris. After single-cell deposition, 10 nL lysis buffer containing 0.1% DDM and 5 mM TCEP in 100 mM TEAB was dispensed into each nanowell. The N2 chip was incubated at 70°C for 45 min in a humidity box to achieve complete cell lysis and protein reduction. Next, 5 nL of 20 mM IAA was added, following by reaction incubation for 30 min in the dark. Proteins were digested to peptide by sequentially adding 0.25-ng Lys-C (5 nL) and 0.5-ng-trypsin (5 nL) into the nanowells and incubating for 3 hours and 8 hours, respectively. For isobaric labeling, we added 50 ng TMT tag in 10 nL ACN into each of the corresponding nanowells according to experimental design. After 1-hr incubation at room temperature, the remaining TMT reagents was quenched by adding 5 nL of 5% HA. Finally, TMT labeled boost (10 ng) and reference (0.5 ng) peptide was distributed into nanowells. The samples were acidified with 5 nL of 5% FA and dried for long-term storage.

### LC-MS/MS analysis

All the samples are analyzed with a nanoPOTS autosampler ^6^ equipped with a C18 SPE column (100 µm i.d., 4 cm, 300 Å C18 material, Phenomenex) and an LC column (50 µm i.d., 25 cm long, 1.7 µm, 130 Å, Waters) heated at 50°C using AgileSleeve column heater (Analytical Sales and Services Inc., Flanders, NJ). Dried samples from nanowell chips or N2 chips were dissolved with Buffer A (0.1% FA in water), then trapped on the SPE column for 5 min. Samples were eluted out from the column using a 120-min gradient from 8% to 45% Buffer B (0.1% FA in ACN) and a 100 nL/min flow rate.

An Orbitrap Eclipse Tribrid MS (Thermo Scientific) operated in data-dependent acquisition mode was employed for all analyses for peptides. Peptides were ionized by applying a voltage of 2,200 V and collected into an ion transfer tube at 200°C. Precursor ions from 400-1800 m/z were scanned at 120,000 resolution with an ion injection time (IT) of 118 ms and an AGC target of 1E6. During a cycle time of 3 s, precursor ions with >+2 charges and > 2E4 intensities were isolated with a window of 0.7 m/z, an AGC target of 1E6, and an IT of 246 ms. The isolated ions were fragmented by high energy dissociation (HCD) level of 34%, and fragments were scanned in an Orbitrap at 120,000 resolution.

### Database searching

All the raw files from the Thermo MS were processed by MaxQuant ^21^ (Ver. 1.6.14.0) with the *UniProtKB* protein sequence database of *Mus musculus* species (downloaded on 05/19/2020 containing 17,037 reviewed protein sequences). Reporter ion MS2 was set as the search type and TMT channel correction factors from the vendor were applied. The mass tolerance for precursor ions and fragment ions was using the default value in MaxQuant. Protein acetylation in N-terminal and oxidation at methionine were chosen as variable modifications, and protein carbamidomethylation in cysteine residues was set as fixed modification. Both peptides and proteins were filtered with a false discovery rate (FDR) of 1% to ensure identification confidence.

### Single-cell proteomics data analysis

The corrected reporter ion intensities from MaxQuant were imported into Perseus (Ver. 1.6.14.0) ^22^ and were log2-transformed after filtering out the reverse and contaminant proteins. Proteins containing >70% valid values in each cell type were considered as quantifiable proteins, and the report ion intensities of the quantified proteins were normalized via the quantile-normalization followed by replacing the missing values based on a standard distribution of the valid values (width: 0.3, downshift: 1.8). To minimize the batch effect from multiple TMT sets, we adjusted the batch effects using the Combat algorithm ^23^ in SVA package, which is embedded in Perseus. Next, the data matrix was separated by cell types and grouped by TMT channel. Combat algorithm is applied to minimize the TMT channel effect. The combined matrix was further applied for statistical analysis, including principal component analysis (PCA) and heatmap hierarchical clustering analysis. ANOVA test was performed to determine the proteins showing statistically significant differences across the three cell types (Permutation-based *FDR* < 0.05, S_0_ = 1), and a 2-way student t-test was applied to explain the significant differences between two groups (*p-value* < 0.05). The processed data were visualized with Graphpad and Perseus.

Proteins intensities without missing values in each cell type in intra-batch or inter-batches were used to calculate the coefficient of variations (CVs). Briefly, for intra-batches, the CVs were calculated using raw protein intensities inside each TMT set and then pooled together to generate the box plots. For inter-batches without batch correction, the CVs were calculated using raw protein intensities across all the TMT sets. To calculate the CVs of intra-batches with batch corrections, raw protein intensities were log2 transformed and followed by imputing missing values. After normalization and batch correction using Combat algorithm ^23^, proteins with imputed values were replaced to ‘NaN’ and filtered out. The protein intensities were exponentially transformed to calculate the CVs.

## Results

### Design and operation of the N2 chip

The N2 chip is distinct from previous nanoPOTS chips ^6, 11, 15, 16^. We cluster an array of nanowells in high density and use each cluster for one multiplexed TMT experiment. In this proof-of-concept study, we designed 9 (3×3) nanowells in each cluster and 27 (3×9) clusters, resulting in total 243 nanowells on one chip (Figure 1a and S1a). Additionally, we designed a hydrophilic ring surrounding the nested nanowells to confine the droplet position and facilitate the TMT pooling and retrieval steps. Compared with previous nanoPOTS chips ^6, 8, 15^, we reduced the nanowell diameters from 1.2 mm to 0.5 mm, corresponding to an 82% decrease in contact areas and an 85% decrease in total processing volumes (Table 1). The miniaturized volume could result in a ∼45× increase in trypsin digestion kinetics because both trypsin and protein concentrations are increased by 6.67×. Both the reduced contact area and increased digestion kinetics are expected to enhance scProteomics sensitivity and reproducibility.

**Table 1.**
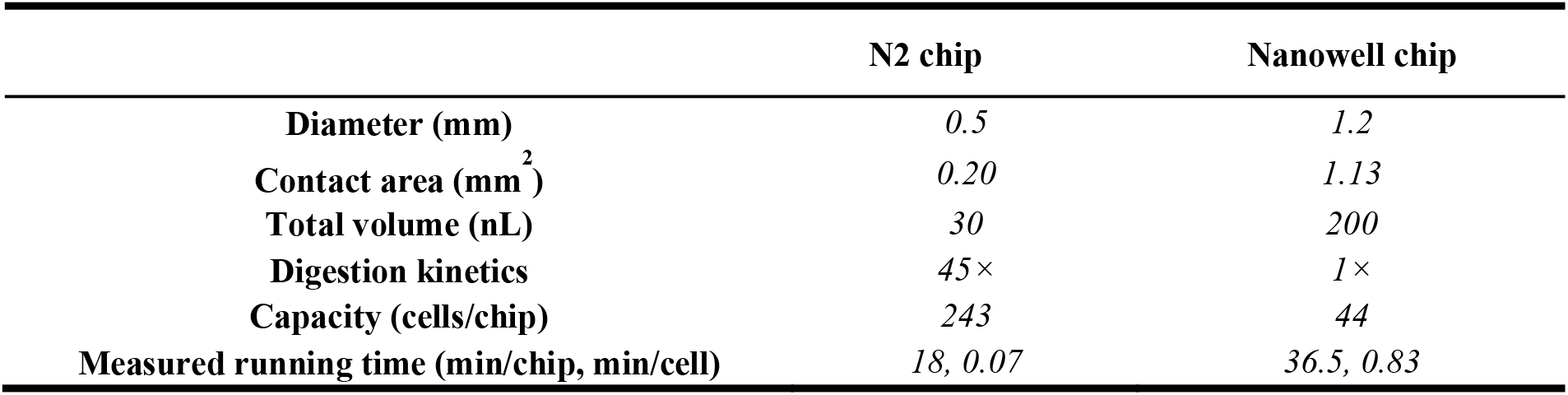
Comparison of technical characteristics between N2 and nanowell chips

|  | N2 chip | Nanowell chip |
| --- | --- | --- |
| Diameter (mm) | 0.5 | 1.2 |
| Contact area (mm<sup>2</sup>) | 0.20 | 1.13 |
| Total volume (nL) | 30 | 200 |
| Digestion kinetics | 45× | 1× |
| Capacity (cells/chip) | 243 | 44 |
| Measured running time (min/chip, min/cell) | 18, 0.07 | 36.5, 0.83 |

The scProteomics sample preparation workflow using the N2 chip is illustrated in Figure 1b. To sort single cells in the miniaturized nanowells, we employed an image-based single-cell isolation system (IBSCI, cellenONE F1.4). The cellenONE system also allowed us to dispense low nanoliter reagents for cell lysis, protein reduction, alkylation, and digestion. After protein digestion, TMT reagent is dispensed to label peptides in each nanowell uniquely. Finally, we distributed 10 ng boosting/carrier peptide and 0.5 ng reference peptide in each nanowell cluster to improve the protein identification rate (Figure S1b) ^14^. To integrate the N2 chip in our LC-MS workflow, we loaded the chip in a nanoPOTS autosampler ^6^. We applied a 3-µL droplet on the rested nanowells, combined the TMT set, and extracted the peptide mixture for LC-MS analysis (Figure 1b). Compared with our previous nanoPOTS-TMT workflow ^6, 15, 16^, the total processing time of each chip was reduced from 36.5 min to 18 min (Figure S1c), which is equivalent to the reduced time from 0.83 min to 0.07 min for each single cell. As such, the N2 chip increases the single-cell processing throughput by >10×.

It is should be noted that the N2 chip can be directly coupled with conventional LC system without the use of the customized nanoPOTS autosampler. As shown in Figure S1d, the user can simply add a 8-µL droplet on the chip and aspirate it back into an autosampler vial for LC injection.

### Sensitivity and reproducibility of the N2 chip

We first benchmarked the performance of N2 chip with our previous nanowell chip using diluted peptide samples from three murine cell lines (C10, Raw, SVEC). To mimic the scProteomics sample preparation process, we loaded 0.1-ng peptide in each nanowell of both N2 and nanowell chips (Figure S1b) and then incubated the chips at room temperature for 2 h. The long-time incubation would allow peptides to absorb on nanowell surfaces and lead to differential sample recoveries. The combined TMT samples were analyzed by the same LC-MS system. When containing at least 1 valid reporter ion value was considered as identified peptides, an average of 5706 peptides was identified with N2 chip, while only 4614 were achieved with nanowell chip (Figure 2a). The increased peptide identifications result in a 15% improvement in proteome coverage; the average proteome identification number was increased from 1082 ± 22 using nanowell chips to 1246 ± 6 using N2 chips (Figure 2b). Indeed, we observed significant increases in protein intensities with N2 chips. The median log2-transformed protein intensities are 13.21 and 11.49 for N2 and nanowell chips, respectively, corresponding to ∼230% overall improvement in protein recovery (Figure 2b). Together, these results demonstrated the N2 chips can dramatically improve the sample recovery and proteomics sensitivity.

**Figure 2.**
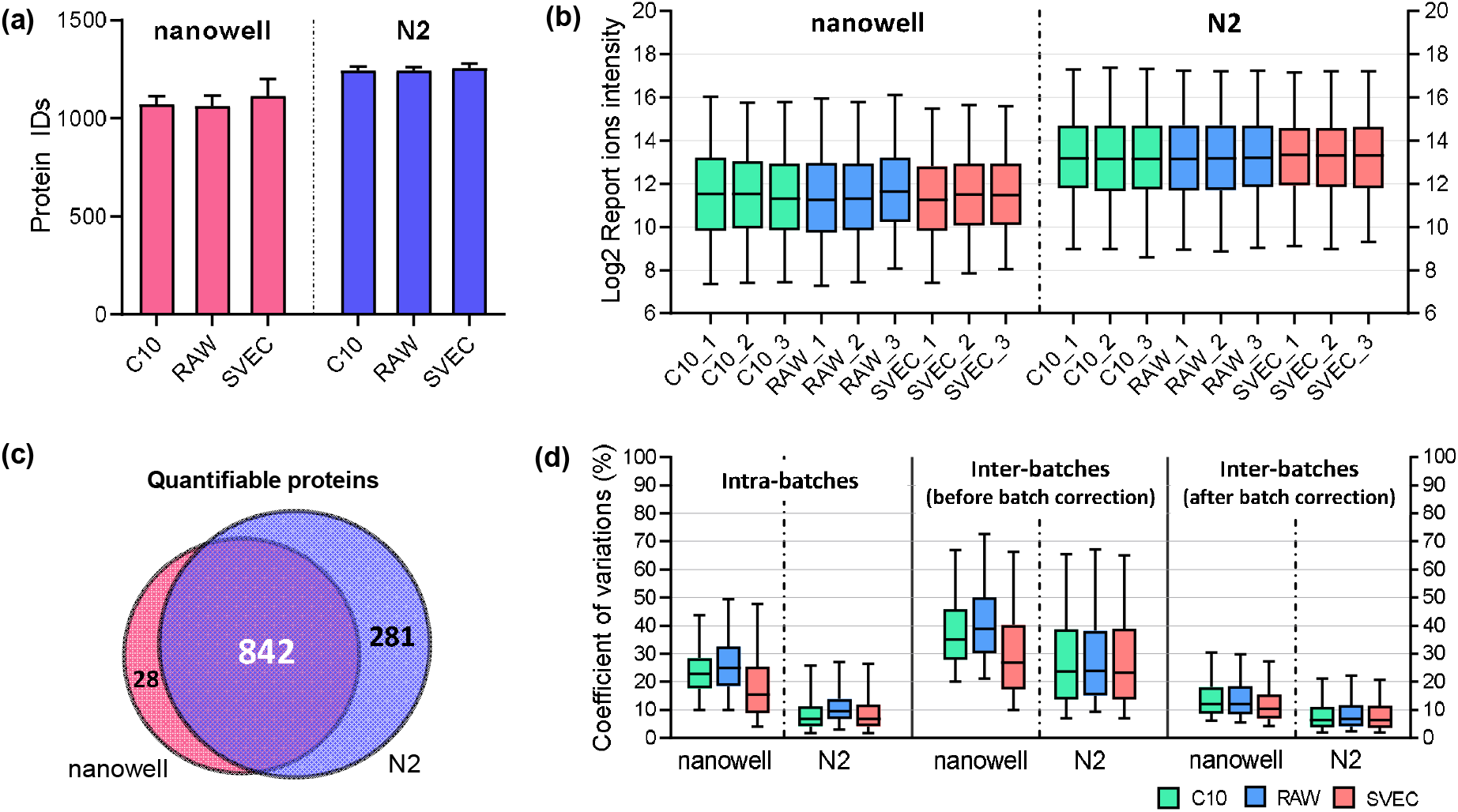
(a) The numbers of protein identifications using nanowell and N2 chips. The error bars indicate standard deviations of 12 samples each containing 0.1 ng tryptic peptides from three cell lines. (b) The distributions of log2 transformed protein intensities in each TMT channel. (c) Venn diagram of quantifiable proteins between nanowell and N2 chips. (d) The distributions of the coefficient of variations (CVs) for proteins identified in each cell type. Protein CVs were calculated inside single TMT batches (left), among different TMT batches without batch corrections (middle), and with batch correction (right).

We assessed if the N2 chip could provide comparable or better quantitative performance compared with nanowell chips. As expected, more proteins are quantifiable with N2 chip if 70% valid values in each cell line were required; the quantifiable protein numbers were 870 and 1123 for nanowell and N2 chips, respectively (Figure 2c). For nanowell chips, pairwise analysis of any two samples showed Pearson’s correlation coefficients from 0.97 to 0.99 between the same cell types and from 0.87 to 0.95 between different cell types (Supplementary figure 2a and 2b). When N2 chips were used, Pearson’s correlation coefficients were increased to a range of 0.98-0.99 for the same cell types, and a range of 0.91-0.96 for different cell types. We next evaluated the quantification reproducibility by measuring the coefficient of variations (CV) of samples from the same cell types. In intra-batch calculations, we obtained median protein CVs of <9.6% from N2 chips, which were dramatically lower than that from nanowell chips (median CVs of < 24.9%) (Figure 2d). Higher CVs were obtained between different TMT batches, which was known as TMT batch effect ^24^. When Combat algorithm ^23^ was applied to remove the batch effect, the median protein CVs from N2 chip dropped to < 6.7%. Such low CVs are similar or even better than other bulk-scale TMT data, demonstrating the N2 chip could provide high reproducibility for robust protein quantification in single cells.

### Proteome coverage of single cells with the N2 chip

We analyzed total 108 single cells (12 TMT sets) from three murine cell lines, including epithelial cells (C10), immune cells (Raw264.7), and endothelial cells (SVEC) (Figure 3a). Noteworthily, these three cell types have different sizes, which allows us to evaluate if the workflow presents a bias in protein identification or quantification based on cell sizes. Specifically, Raw cells have a diameter of 8 µm, SVEC of 15 µm and C10 of 20 µm. (Figure S3a). A total of 108 individual cells were analyzed, corresponding to 12 separate multiplexed TMT sets.

**Figure 3.**
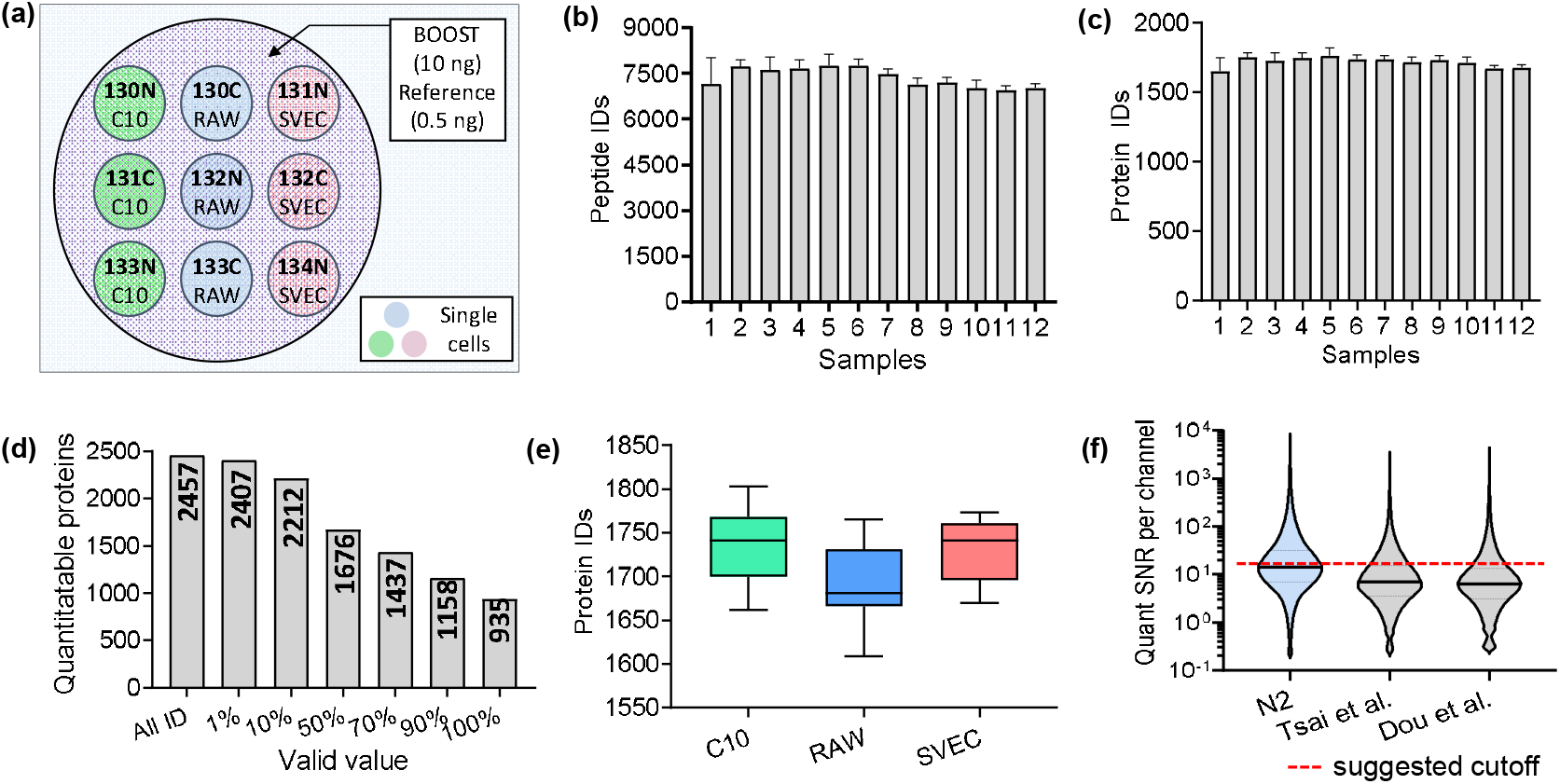
(a) Experiment design showing single-cell isolation and TMT labeling on the N2 chip. (b, c) The average numbers of identified peptides and proteins for the 12 TMT sets. At least 1 valid value in the 9 single-cell channels is required to count as an identification. Error bars show the standard deviations from 9 single cells. (d) The numbers of quantifiable proteins based on different percentages of required valid values. (e) Box plot showing the distributions of protein identification numbers. (f) Violin plot showing the distributions of signal to noise ratio (SNR) per channel for raw single-cell signals calculated by SCPCompanion. ^17^ Two published TMT scProteomics datasets from our group using nanowell chip ^15, 16^ were used to benchmark the data generated in this study.

Among the 12 TMT sets, our platform identified an average of ∼7369 unique peptides and ∼1716 proteins from each set with at least 1 valid value in the 9 single-cell channels (Figure 3b and 3c). We identified total 2457 proteins, and 2407 proteins had reporter ion intensities in at least 1 single cells across the 108 cells (Figure 3d). When a stringent criteria of >70% valid values was applied, the number of proteins dropped to 1437. As expected, we observed the numbers of proteins identified for three cell types ranked according to the cell sizes (Figure S3a). Average 1735, 1690, and 1725 proteins were identified in C10, RAW, and SVEC cells, respectively (Figure 3e). In addition, similar trends were also observed in the distribution of protein intensities (Figure S3b).

Cheung and coworkers ^17^ recently introduced the software SCPCompanion to characterize the quality of the data generated from single-cell proteomics experiments employing isobaric stable isotope labels and a carrier proteome. SCPCompanion extracts signal-to-noise ratio (SNR) of single-cell channels and provides cutoff values to filter out low-quality spectra to obtain high-quality protein quantitation. In line with our experimental design, SCPCompanion estimated ∼0.1 ng proteins were contained in single cells and the boost-to-single ratio is ∼100 (Supplementary Table 1), indicating minimal peptide losses in the N2 chip. More importantly, the median SNR per single-cell sample was 14.4, which is very close to the suggested cutoff value of 15.5, corresponding to ∼50% of raw MS/MS spectra can provide robust quantification. We also compared the data quality generated with previous nanowell chips and similar LC-MS setup ^15, 16^. The median SNR values per sample were 7.0 ^15^ and 6.4 ^16^, which were significantly lower than the data generated by the N2 chip (Figure 3f).

### Cell typing with scProteomics

To assess the quantitative performance of the N2 chip-based scProteomics platform, we first performed a pair-wise correlation analysis using the 1437 proteins across the 108 single cells. As expected, higher correlations were observed among the same types of cells and lower correlations among different types of cells (Figure 4a). The median Pearson correlation coefficients are 0.98, 0.97, and 0.97 for C10, RAW, and SVEC cells, respectively. We next calculated the coefficient of variations (CVs) using protein abundances for the three cell populations. Interestingly, we see very low variations with median CVs < 16.3% (Figure S4), indicating protein expression are very stable for cultured cells under identical condition. Principal component analysis (PCA) showed strong clustering of single cells based on cell types and the three clusters were well separated from one another (Figure 4b). We compared the our previous PCA result for the same three cell types using nanowell-based platform (Figure S5) ^16^. The N2 chip not only increased the proteome coverages, but also dramatically improved the classification power.

**Figure 4.**
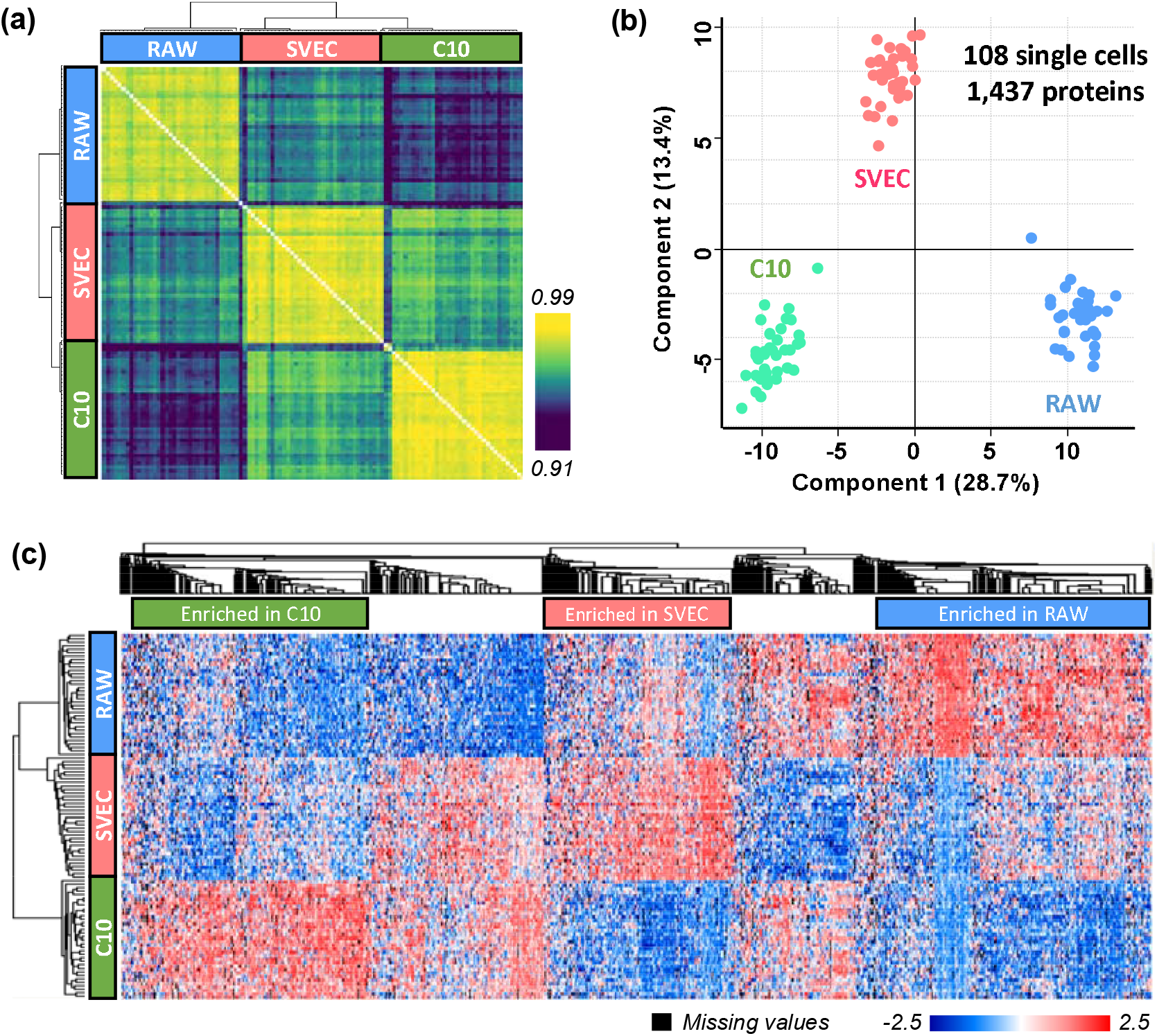
(a) Clustering matrix showing Pearson correlations across 108 single cells using log2-transformed protein intensities, and (b) PCA plot showing the clustering of single cells by cell types. Total 1437 proteins were used in the PCA projection. (c) Heatmap with hierarchical clustering showing 1,127 significant proteins based on ANOVA test. Three protein clusters used for pathway analysis were labeled and highlighted.

To identify proteins leading the clustering of the three cell populations, an ANOVA test was performed (Permutation-based FDR < 0.05, S0 = 1). Of the total 1437 proteins, 1127 proteins were significantly differentially changed across three cell types (Figure 4c). Among them, 237 proteins were enriched in C10 cells, 203 proteins were enriched in SVEC cells, and 275 proteins were enriched in RAW cells.

Proteins enriched in each cell type revealed differences in molecular pathways based on the REACTOME pathway analysis (Figure S6). For example, the proteins higher in abundance in C10 cells were significantly enriched in REACTOME terms such as “vesicle-mediated transport”, “membrane trafficking”, “innate immune system”, or “antigen processing-cross presentation”. These functions are in line with the known functions of lung epithelial cells, of which the C10 are derived from ^25^. The protein more abundant in RAW cells, which derive from murine bone marrow macrophages, were enriched in REACTOME terms associated with “neutrophil degranulation”, “innate immune system” in line with their immune function. Other REACTOME terms related to the “ribosome” and the “pentose phosphate pathway” were also enriched. These pathways not only suggest that there is intricate cooperation between macrophages and neutrophils to orchestrate resolution of inflammation and immune system ^26^, but also show that system metabolism strongly interconnects with macrophage phenotype and function ^27^. Proteins more abundant in SVEC cells (murine endothelial cells) were enriched in pathways include “processing pre-mRNA”, “cell cycle”, or “G2/m checkpoints”. This suggests its proliferation, migration, or coalescing of the endothelial cells to form primitive vascular labyrinths during angiogenesis ^28^.

### Identifying cell surface markers with scProteomics

One of most unique advantages of scProteomics over single-cell transcriptomics is the capability to identify cell surface protein markers, which can be used to enrich selected cell populations for further studies. We next assessed if we can use our scProteomics data to identify cell-type-specific membrane proteins for the three cell populations. We matched the enriched protein lists to a subcellular-component database on UniProtKB, which consists 2,871 of reviewed plasma membrane proteins for Mus musculus (updated on 01/04/2021). We generated a list containing 63 plasma membrane proteins (Supplementary Table 2). Among them, 16 plasma membrane proteins were highly expressed in C10 compared to RAW and SVEC cells, while 34 and 13 plasma membrane proteins were significantly enriched in RAW and SVEC cells, respectively. For example, EZRI ^29^, JAM1 ^30^, and NCAM1 ^31^, which were previously known to protect the barrier function of respiratory epithelial cells by enhancing the cell-cell adhesion, were highly expressed in C10 cells (Figure 5, left panel). For RAW enriched membrane proteins, CD14 ^32^ and CD68 ^32, 33^ are mainly produced in macrophage cells and widely used as a histochemical or cytochemical marker for inflammation-related macrophages (Figure 5, middle panel). CY24A is a sub-component of the superoxide generating NOX2 enzyme on macrophage membrane ^34^. In term of SEVC enriched protein markers, BST2 is known to highly express in blood vessels throughout the body as an intrinsic immunity factor (Figure 5, right panel) ^35^. HMGB1 and DDX58, which were found to be highly expressed in endothelial cells in lymph node tissue based on tissue microarray (TMA) results in human protein atlas, could also be used as protein markers to differentiate SVEC cells with other two cell types. We also attempted to compare with our previous results using nanowell chips (Figure S7) ^16^. Only 5 out of the 9 membrane proteins could be classified as cell surface markers and 3 proteins were not detected, likely due to low sensitivity and reproducibility. Together, these data indicate the feasibility of using scProteomics to identify cell-type-specific membrane proteins for antibody-based cell enrichment.

**Figure 5.**
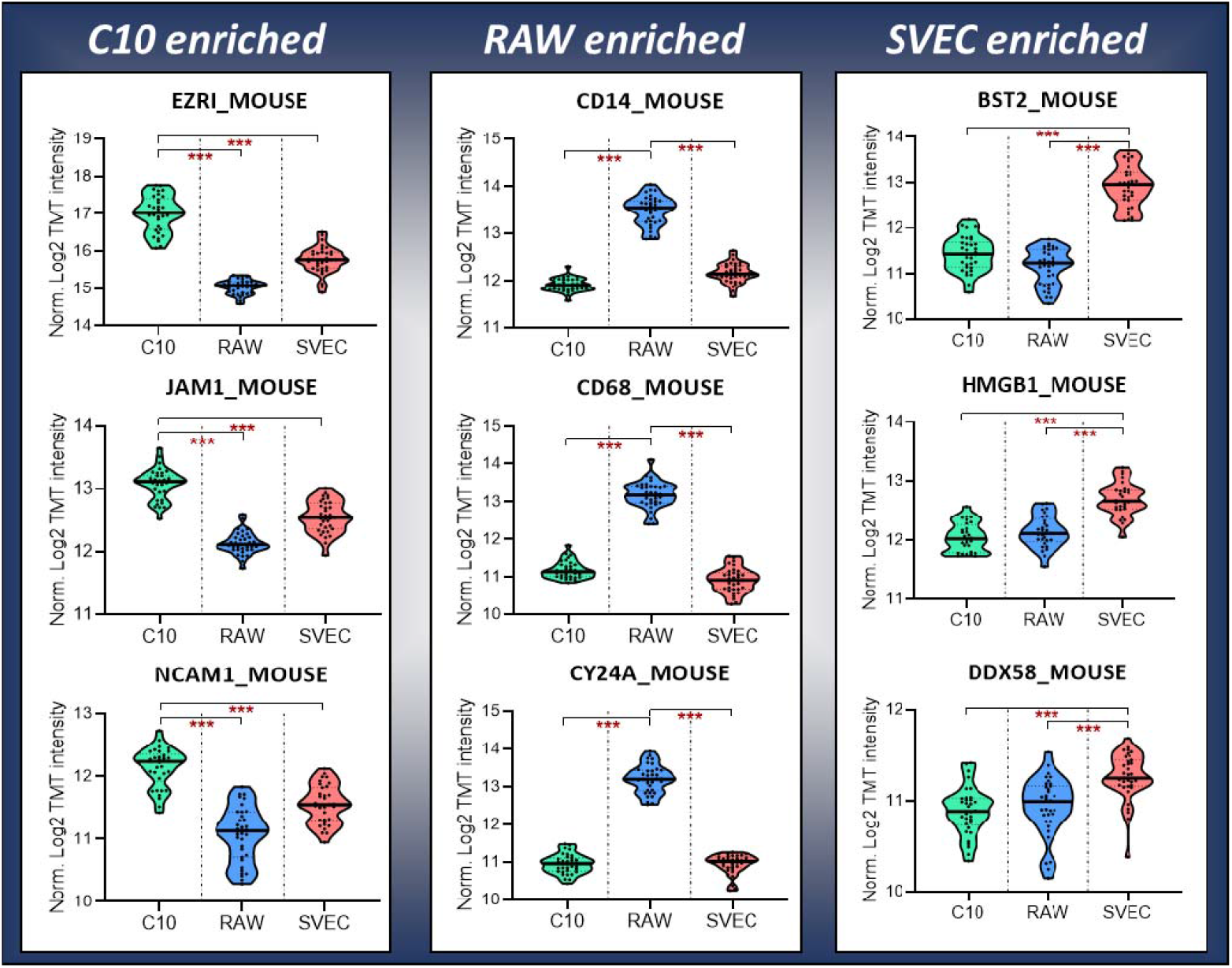
Violin plots showing nine putative plasma membrane proteins enriched in three cell types. Protein in each column are statistically significant (p-value<0.001) expressed in the specific cell type.

## Discussion

We have developed a high-throughput and streamlined scProteomics sample preparation workflow based on nested nanoPOTS array (N2) chips. The N2 chips reduce nanowell volumes to ∼30 nL and improve the protein/peptide sample recovery by 230% compared with our previous nanoPOTS chips ^15, 16^. The N2 design also significantly simplifies the TMT-based isobaric labeling workflow by eliminating the tedious sample pooling step (e.g., aspirating, transferring, and combining). Using the N2 chip, 243 single cells can be analyzed in a single microchip, representing 5× more numbers than our previous chips. In the near future, we envision the development of higher capacity N2 chips and/or stable isotope isobaric labeling reagents to enabling higher multiplexing scProteomics experiments (e.g. over 1000 cells per chip containing 5×5 array and 40 total clusters).

Using a recently-developed software, SCPCompanion^17^, we observed single-cell SNRs were dramatically improved with N2 chip-based scProteomics workflow. The improvement results in high Pearson correlations (median R of ∼ 0.97) of single cells from the same cell lines. Importantly, we observed low protein expression variations (median CVs of ∼16.3%), suggesting the proteome is highly stable for single cells under identical culture conditions. These observations suggest cultured cells are good models to evaluate and benchmark the quantitative performance of scProteomics technologies.

In the analysis of three different cell lines, we verified the scProteomics can robustly classify single cells based on protein abundances and reveal functional differences among them. We also showed it was possible to directly identify cell surface markers by leveraging established subcellular-component databases.

It should be noted that all the single-cell isolation and sample preparation were performed using a commercially available system (cellenONE). The microchip fabrication can be readily implemented in a typical cleanroom facility. Thus, we believe our N2 chip-based scProteomics workflow can be rapidly disseminated.

In summary, we believe the N2 chip provides a universal scProteomics platform with broad applications in studying cell differentiation, tumor heterogeneity, rare cells from clinical specimens.

## Supporting information

SCPCompanionResults

PlasmaProteinMarkerList

## Acknowledgments

We thank Matthew Monroe for helping with data deposition. This work was supported by a Laboratory Directed Research and Development award (Y.Z.) from Pacific Northwest National Laboratory and a Intramural program (Y.Z.) at EMSL (grid.436923.9), a DOE Office of Science User Facility sponsored by the Office of Biological and Environmental Research and operated under Contract No. DE-AC05-76RL01830. Part of this work is also supported by NIH grants U01 HL122703 (C.A.) and P41 GM103493 (R.D.S.).

## Contributions

Y.Z conceptualized and designed the research. S.M.W, A.V.V. and H.S.M. fabricated N2 microchips. J.W., R.L.S, L.M.M., and J.C.B. performed the cell culture and sample preparation. S.M.W. and R.J.M performed LC-MS analysis. J.W., G.C., and Y.Z. analyzed data. J.W., G.C., R.D.S., L.P.T., and Y.Z. wrote the manuscript.

## Data Availability

The mass spectrometry proteomics data have been deposited to the ProteomeXchange Consortium via the MassIVE partner repository with the dataset identifier MSV000086809.

---

**Supplementary figure 1.**
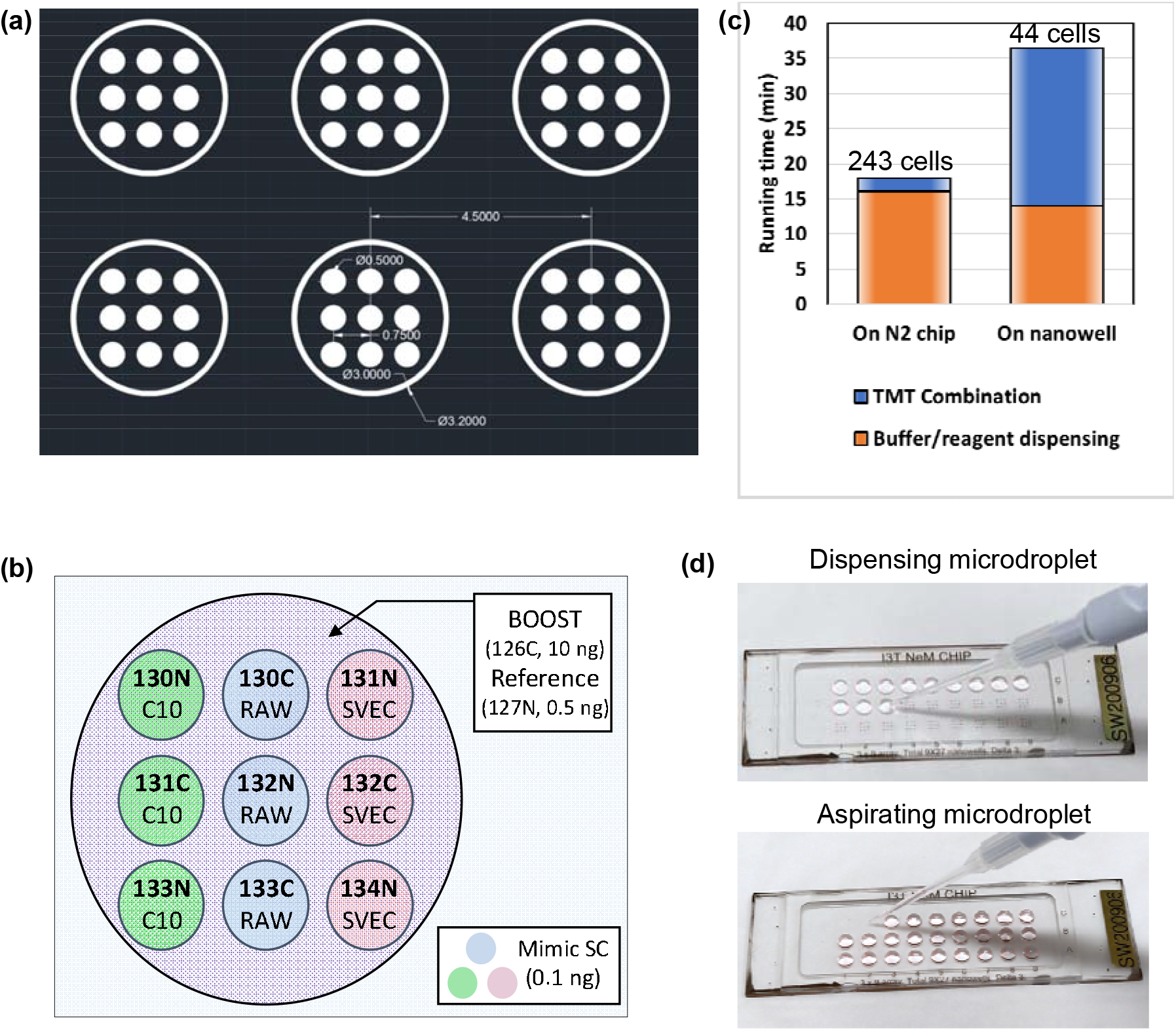
(a) Schematic illustration showing the N2 chip design. The white areas are designed for nanowells and hydrophilic rings, while the rest is hydrophobic surface. (b) Experiment design for mimic single-cell sample (0.1 ng peptide) on the N2 chip. (c) Estimated robot operation time for single cell proteomics using N2 chip and nanowell chip. (d) Photographs showing TMT-based samples can be pooled together by spotting a 6-µL droplet using a micropipette. Similarly, the pooled sample can be retrieved and loaded into an autosampler vial for LC-MS analysis.

**Supplementary figure 2.**
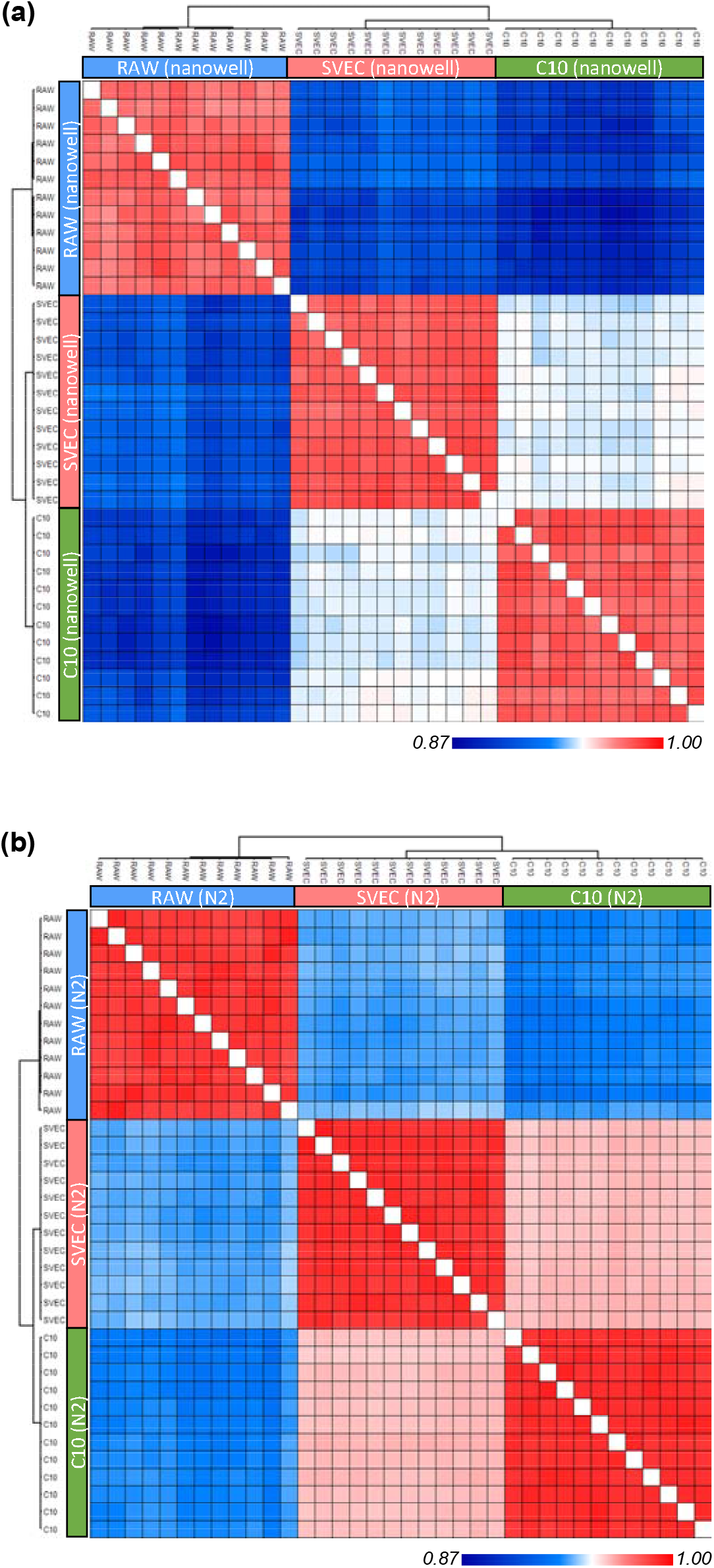
Heatmap of pairwise Pearson correlations among individual samples in nanowell chip (a) and N2 chip (b).

**Supplementary figure 3.**
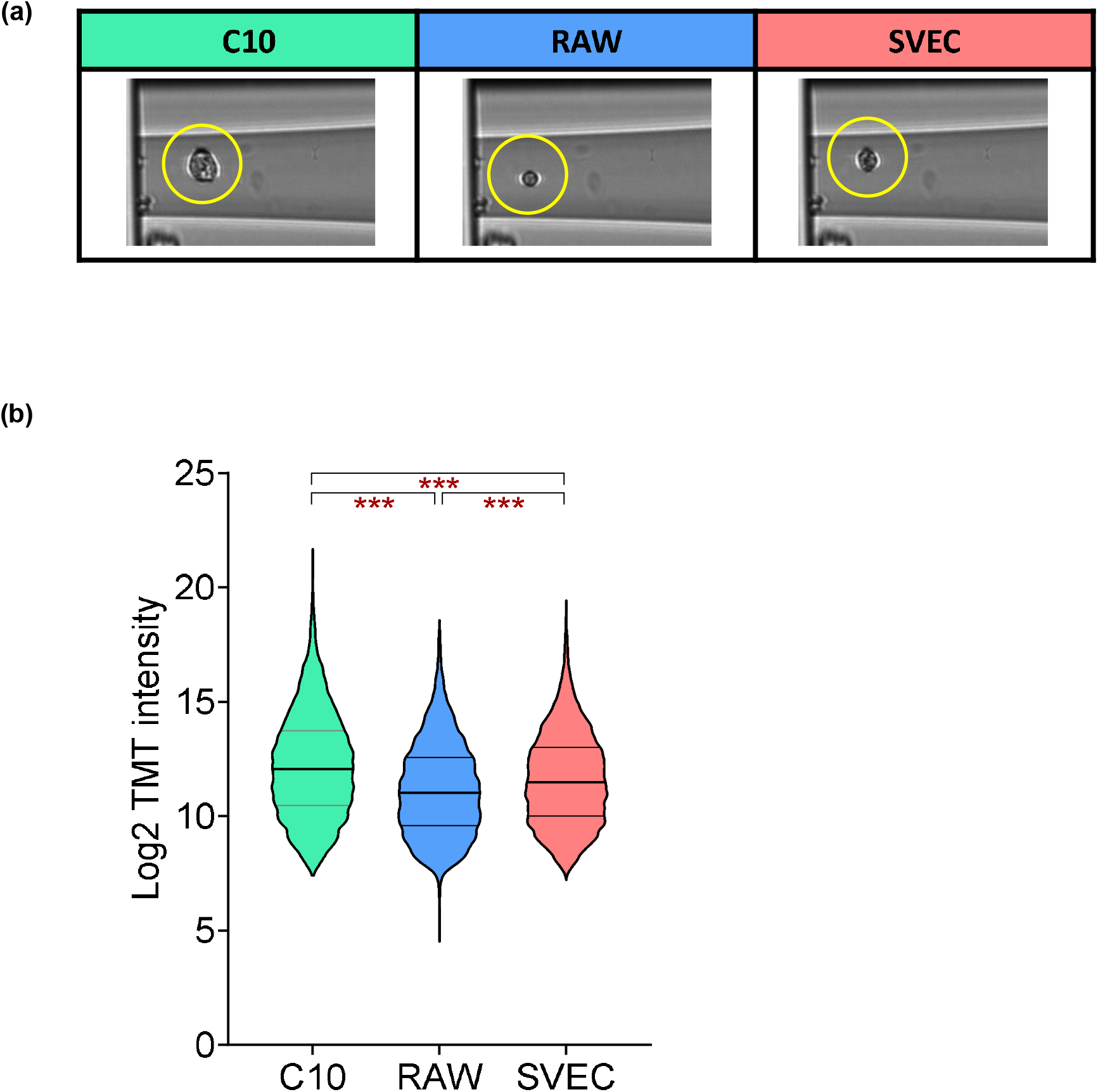
(a) Representative images of single cells. The measured cell sizes in diameter are 18-20 µm for C10 cells, 7-10 µm for RAW cells, and 13-15 µm for SVEC cells. (b) Violin plots showing the distribution of log2 transformed protein intensities for the three cell types.

**Supplementary figure 4.**
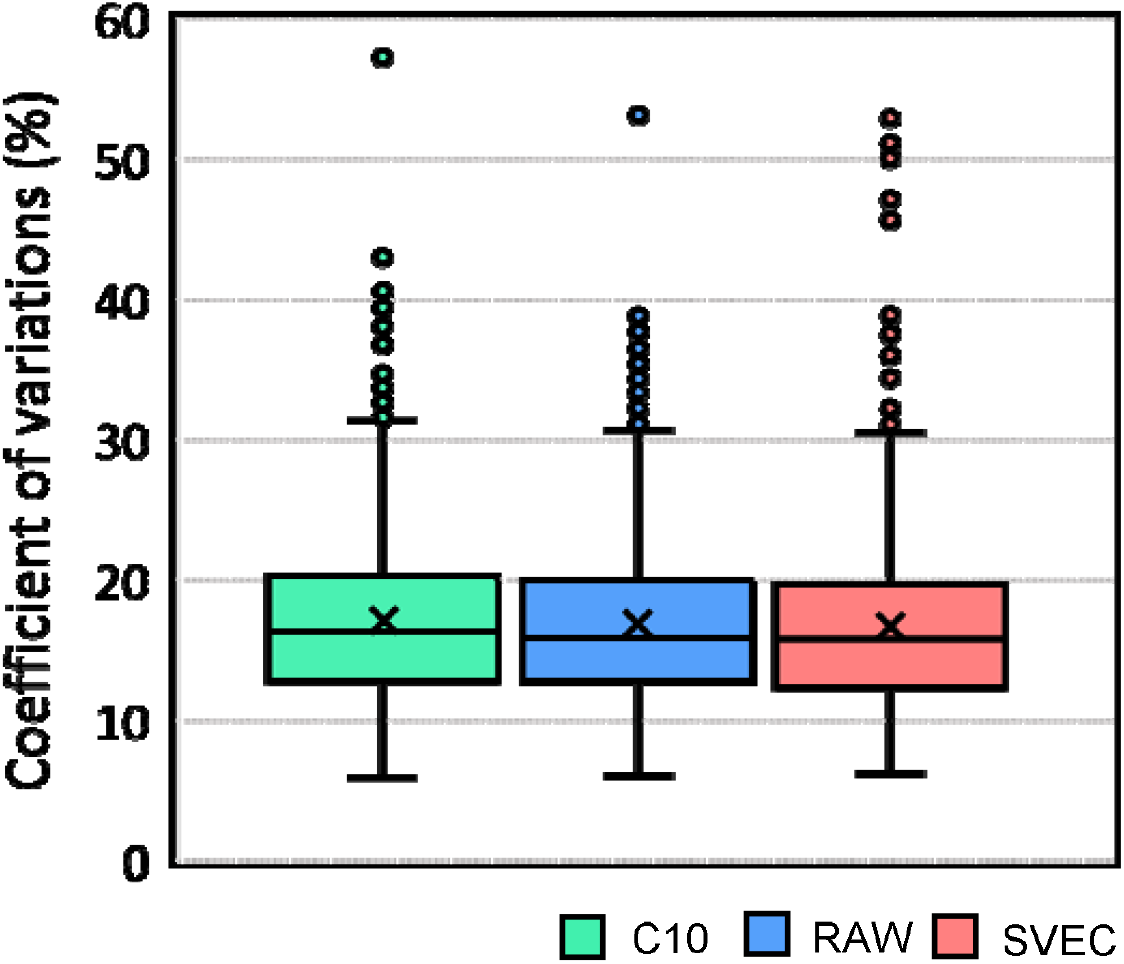
The distribution of coefficient of variations (CVs) for protein abundances in single cells among inter TMT batches with batch correction. For each cell type, 36 single cells from 12 TMT sets were used for the calculation.

**Supplementary figure 5.**
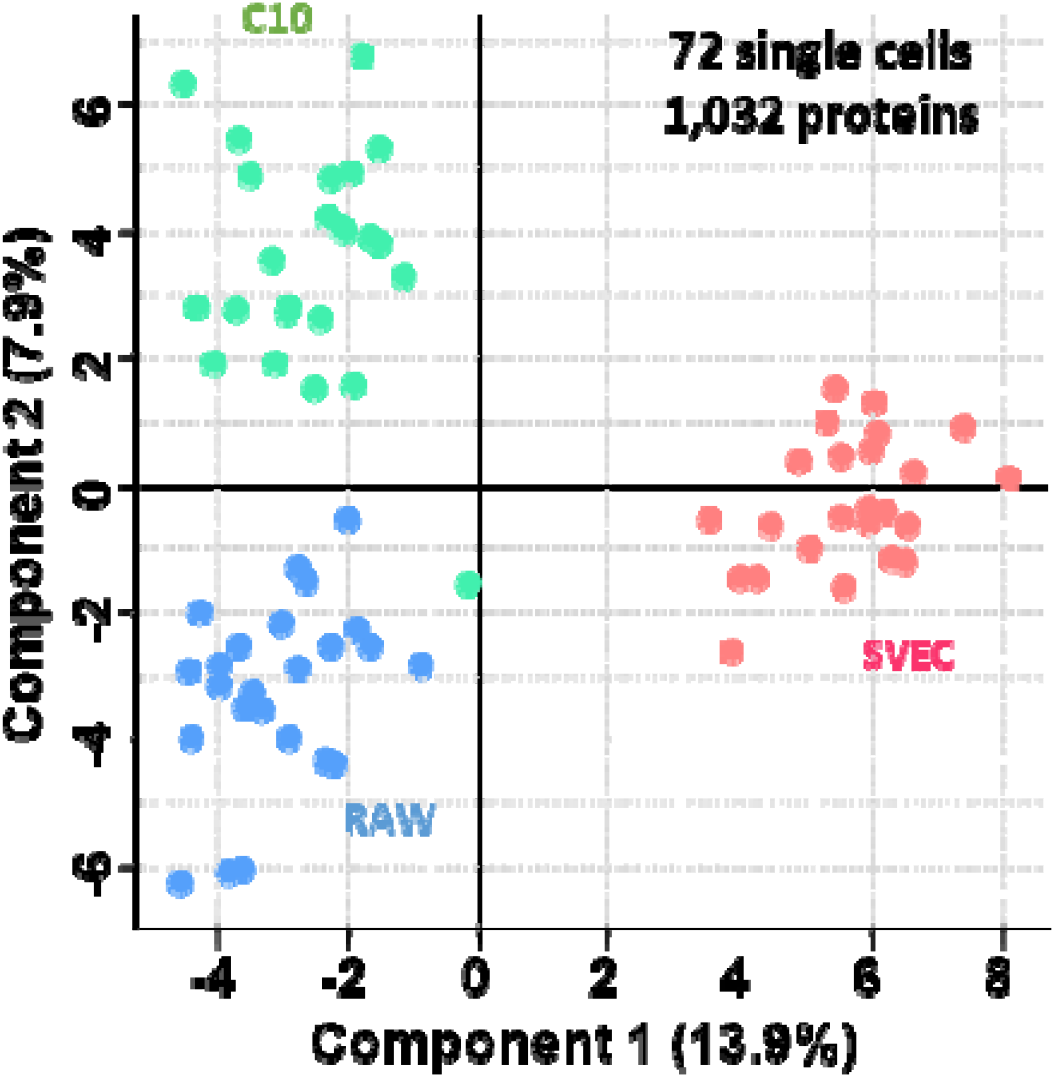
PCA plot showing the clustering of 72 single cells using nanowell chips^16^. Total 1032 proteins were used.

**Supplementary figure 6.**
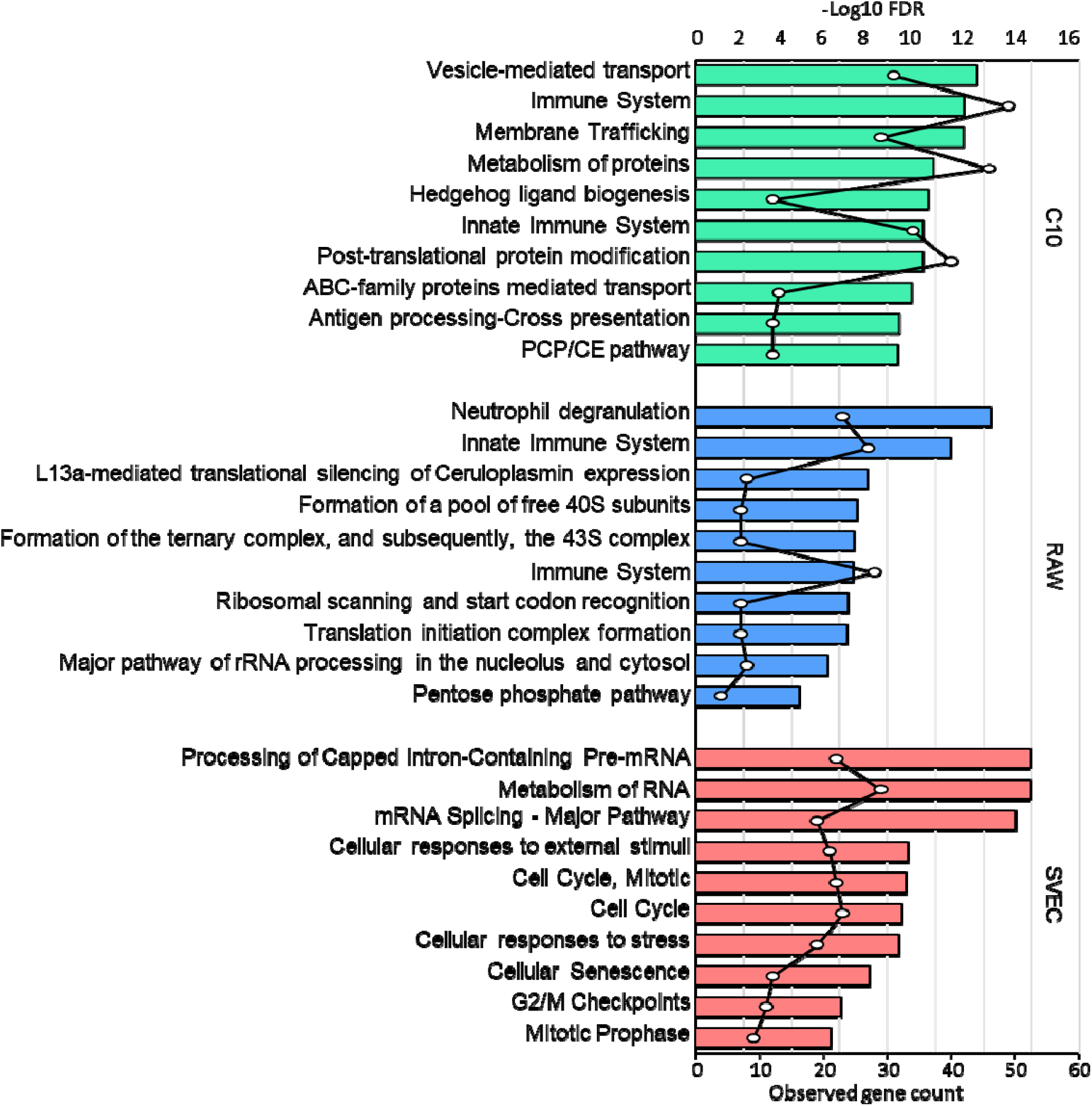
Reactome pathway analysis of enriched proteins in each cell type by hierarchical clustering analysis. The top 10 of the pathways per each homogeneous cell type based on adjusted p-value were listed with the number of observed protein count.

**Supplementary figure 7.**
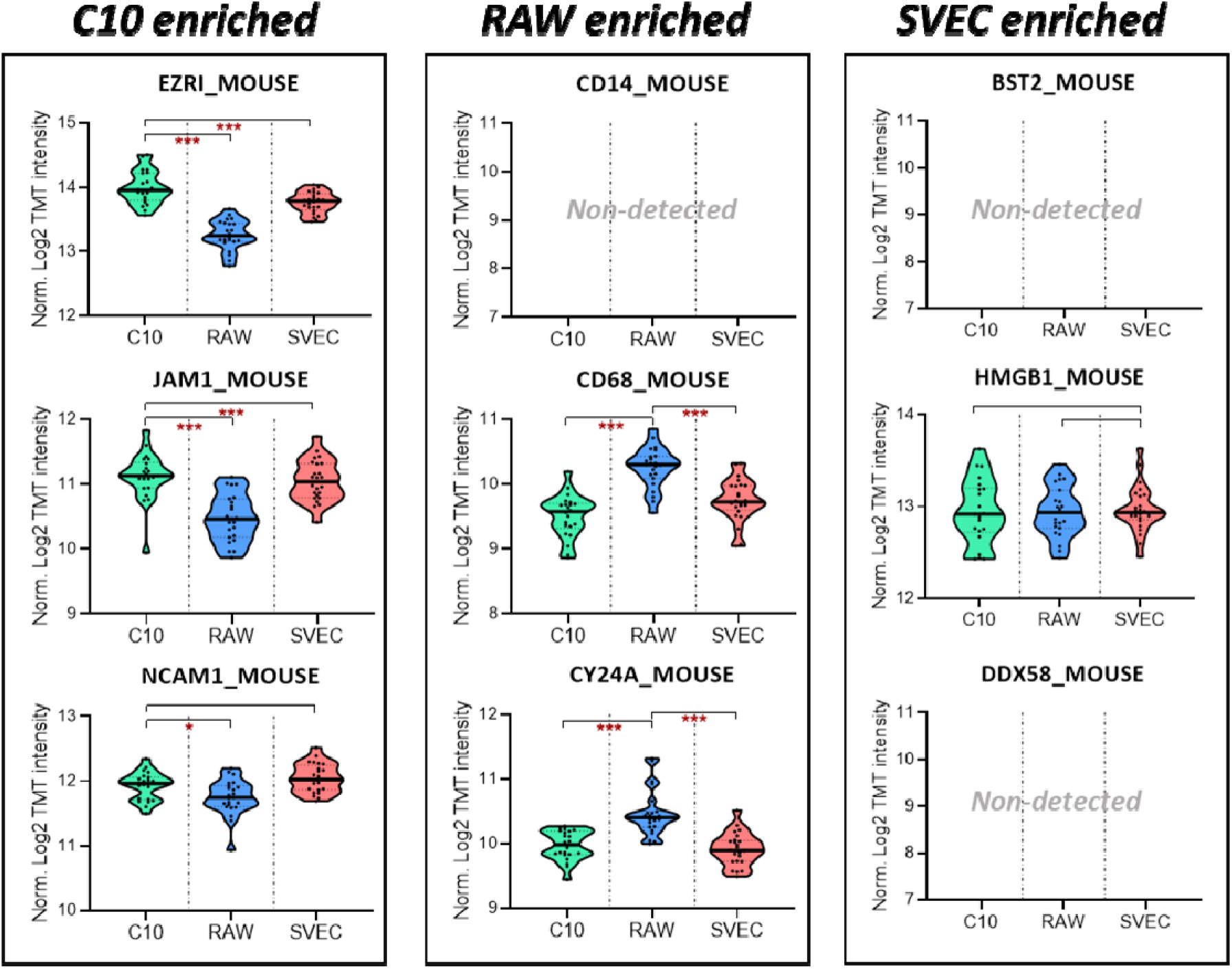
Violin plots showing the intensity distributions of putative plasma membrane protein markers for specific cell types using previous nanowell-chip-based single-cell proteomics data. The protein lists were selected based on the data from the N2 chips and shown here to compare the improved proteome coverage and quantitation performance of the N2 chip.

